# Characterization and targeting of the endosomal signaling of the gastrin releasing peptide receptor in pruritus

**DOI:** 10.1101/2025.03.17.643743

**Authors:** Jeffri Retamal Santibañez, Diana Bok, Shavonne Teng, Divya Bhansali, Marcella de Amorim Ferreira, Raquel Tonello, Chloe J. Peach, Rocco Latorre, Gokul SA Thanigai, Kam W. Leong, Dane D. Jensen

## Abstract

Chronic pruritus is a major unmet clinical problem affecting one in four adults. G protein-coupled receptors (GPCRs) are key receptors driving itch signaling and are a therapeutic target for itch relief. The endosomal signaling of GPCRs provides new challenges for understanding how GPCR signaling is regulated, how endosomal signaling of GPCRs contributes to disease states like chronic pruritus and opens new targets for therapeutic development. The Gastrin releasing peptide receptor (GRPR) is a key mediator of pruritus in the spinal cord. Yet, little is known about the molecular mechanisms regulating GRPR signaling in pruritus, if GRPR can signal from endosomes, or the role of endosomal GRPR in the development of pruritus. Here we show the importance of internalization and endosomal signaling of GRPR in pruritus. Agonist induced GRPR internalization and trafficking was quantified using BRET or microscopy while endosomal-mediated ERK signaling was measured using compartmentalized FRET biosensors. Recruitment of G proteins to endosomes was measured with NanoBit BRET. pH sensitive mesoporous silica nanoparticles (MSN) which accumulated in endosomes were used to deliver RC-3095, a GRPR specific antagonist, intracellularly to block endosomal signaling of GRPR. MSN-RC proved more effective than free RC-3095 at inhibiting chloroquine scratching in mice. Our results demonstrate a critical role for GRPR endosomal signaling in itch sensation. These results highlight the ability of endosomally targeted antagonist to inhibit GRPR signaling and provide a new target for developing therapeutics that block GRPR mediated pruritus.

**Significance Statement:** GPCRs are dynamic signaling receptors that can continue to signal following internalization and trafficking to endosomes. Using subcellular targeted BRET and FRET based biosensors we can quantify the recruitment of signaling partners like G proteins and arrestins to GRPR from the endosomal compartment. Inhibition of clathrin and dynamin mediated endocytosis allowed to differentiate plasma membrane vs endosomal signaling of GRPR. pH sensitive nanoparticles loaded with the GRPR antagonist RC-3095 are endocytosed to the endosomal network where they specifically target and block endosomal GRPR signaling. Intrathecal injection of RC-3095 loaded nanoparticles blocked chloroquine induced scratching behavior in mice. Thus, intracellular GRPR drives itch sensation and targeted inhibition of intracellular GRPR signaling is a more effective strategy to treat pruritus.

## Introduction

Itch is a sensory modality that is defined as an unpleasant sensation that elicits the desire to scratch. Itch has evolved to protect organisms from harmful agents and is conveyed through pruritogenic sensory pathways. The presence of the pruritogens are detected by heterotrimeric G protein-coupled receptors (GPCRs) expressed on primary sensory neurons (1). GPCRs are heterotrimeric transmembrane proteins and are the largest receptor family. GPCRs are activated by a host of agonists and due to their ubiquitous expression, are involved in most physiological processes including nociception and pruritus. The importance of GPCRs in sensing pruritogens in the periphery and in transmitting itch sensation in the central nervous system continues to evolve (2, 3). GPCRs participate in itch transmission at multiple levels. GPCRs on primary sensory neurons provide the principal mechanism for detection of pruritogens (1). Pruritogen-sensing GPCRs include the large family of Mas-related G protein-coupled receptors (Mrgprs including, but not limited to, A3, C11, X1, and X4), as well as receptors for histamine, serotonin, bile acids, neuropeptides, proteases, and cytokines (4–16). Pruritogenic GPCRs on primary sensory neurons often generate signals that activate or sensitize members of the transient receptor potential (TRP) family of ion channels, notably TRP ankyrin-1 (TRPA1)(17–19). These processes initiate neuronal excitation and central transmission of itch. Pruriceptive sensory neurons that transmit itch release glutamate (L-Glut), natriuretic polypeptide B (NppB), and gastrin releasing peptide (GRP) which in turn activate excitatory spinal interneurons expressing the gastrin releasing peptide receptor (GRPR) which transmit itch sensation to cortical projection neurons (2, 20–24). GRPR is a GPCR expressed on excitatory spinal interneurons in lamina I of the spinal cord that receive input from sensory neurons (25). GRPR is a final common path for non-histaminergic itch as previous studies have shown intrathecal injections of GRPR antagonists block scratching behavior in mice (25). Additionally, ablation studies of GRPR positive interneurons confirmed GRPR positive interneurons regulate scratching responses to pruritogens in mice (25, 26). The mechanisms regulating the activation of these interneurons are of key interest because pruriceptive activation tends to involve a prolonged “waxing and waning” cycle (27) where onset of activity is slow but builds to a persistent state that remains elevated even after the pruritogen has been removed. This slow build was characterized in GRPR interneurons where activation and depolarization required prolonged volumetric release of GRP and the resultant prolonged activation of GRPR (20). This prolonged release of GRP would result in the increased trafficking of GRPR to the endosomal pathway in interneurons. Endosomal signaling of GRPR would be one mechanism to allow the building up and prolonged activation of these excitatory interneurons.

GRPR activation, like many GPCRs, is tightly regulated as GPCRs are dynamic signaling proteins that can adapt multiple conformations. GRPR is a Gα_q_ coupled GPCR and after activation by binding GRP, is rapidly phosphorylated by G protein kinase 2 (GRK2), bound by βarrestins (βARR) and internalized (28–31). Traditionally, GPCR signaling was proposed to be confined to the plasma membrane where activated GPCRs activated intracellular G proteins and then were rapidly phosphorylated and endocytosed to arrest signaling (31, 32). We now understand that the endocytosis of GPCRs does not terminate receptor signaling but instead provides a mechanism to shift signaling cascades from the plasma membrane to intracellular locations. GPCRs can recruit signaling partners like βARRs to form a “signalosome” at intracellular compartments and continue to signal *via* G protein-dependent and independent mechanisms (33–38). Previous studies have highlighted the importance of intracellular GPCR signaling in prolonged neuronal activation in chronic pain and inflammation (33, 39). Pagani *et al.* defined a novel mechanism whereby GRPR signaling regulated itch via slow but prolonged activation of GRPR through repeated release of GRP (20). Since GRPR was reported to undergo rapid desensitization and internalization at the plasma membrane (28, 31), we hypothesized that endosomal signaling of GRPR would be one mechanism to prolong GRPR activation and regulate the spinal gate controlling itch. Herein, we demonstrate GRPR can recruit G proteins to the endosomal compartment and that inhibiting GRPR internalization using inhibitors of clathrin or dynamin block GRPR internalization, inhibiting scratching responses in mice. Moreover, using pH sensitive nanoparticles designed to release their cargo, GRPR antagonist, in the acidic environment of the endosomes, results in a greater inhibition of scratching behavior than injection of the free drug.

## Results

### Endocytosis and trafficking of GRPR to endosomes

Stimulation of GRPR with GRP results in the endocytosis of GRPR to intracellular compartments. Here we used confocal microscopy and Bioluminescence Resonance Energy Transfer (BRET) assays to visualize and quantify the trafficking of GRPR. HEK cells expressing the fluorescent tagged GRPR-mApple and the early endosome CellLight were challenged with GRP (10 nM). GRPR-mApple moved from the plasma membrane and colocalized with the early endosomal marker Rab5a-GFP (**Fig. 1 A**). GRPR-mApple endocytosis was shown to be clathrin and dynamin dependent as preincubation with the dynamin inhibitor Dyngo4a (Dy4, 30 µM) and the clathrin inhibitor PitStop2 (PS2, 30 µM) attenuated GRPR-mApple release from the plasma membrane and internalization to early endosomes. Inactive analogues of PS2 (PS2Inact, 30 µM) and Dy4 (Dy4Inact, 30 µM) had no impact on GRPR-mApple trafficking to early endosomes (**Fig. 1 A**).

**Figure 1.**
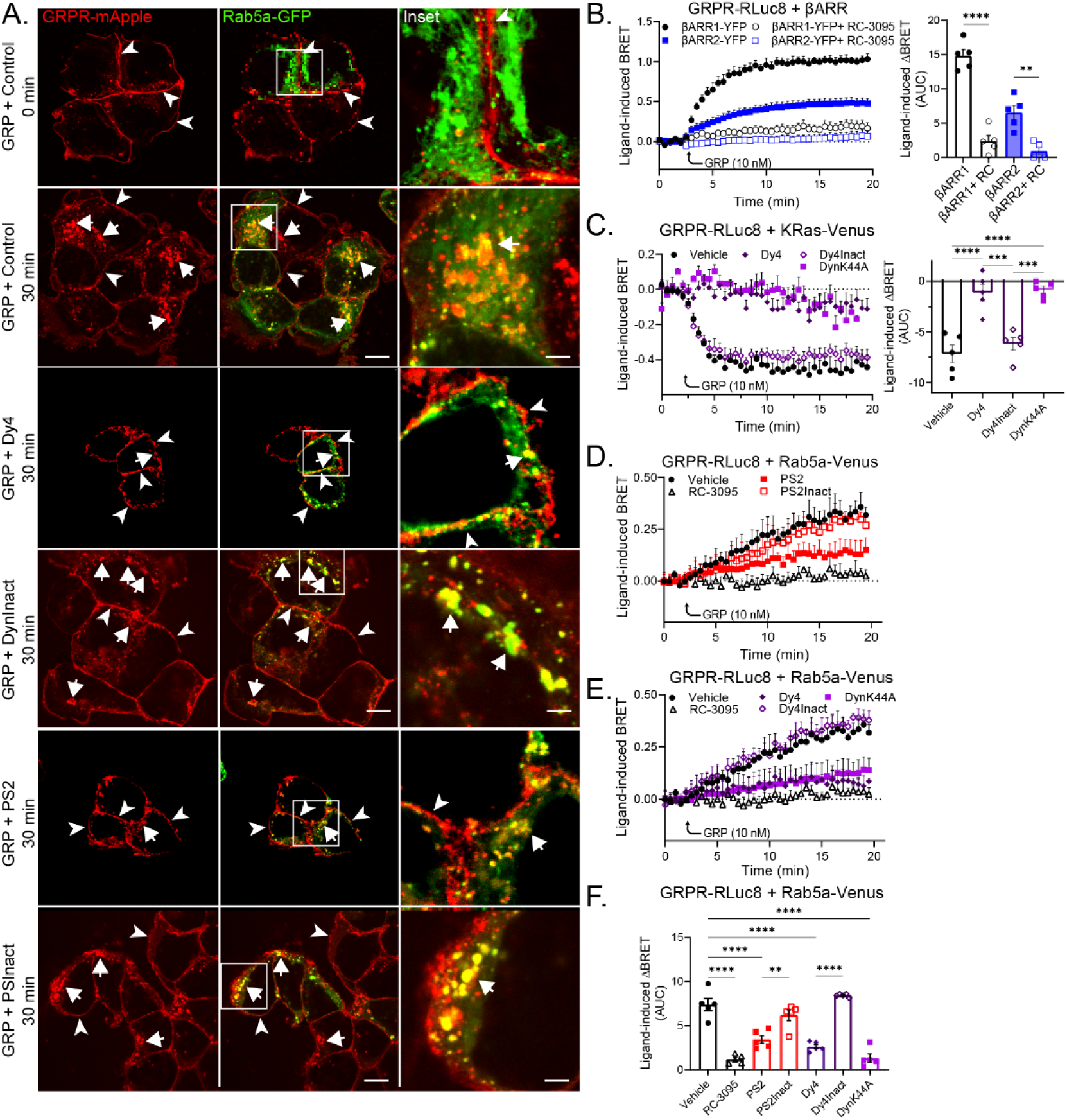
GRPR endocytosis can be blocked by clathrin and dynamin inhibitors. **A**. Plasma membrane bound GRPR-mApple (Arrow heads) is endocytosed to Rab5a-GFP positive endosomes at 30 min following activation by GRP (Arrows) in HEK293 cells pretreated with vehicle, Dy4 Dy4Inact, PS2, or PS2Inact. Scale 10 µm, inset scale is 2.5 µm, representative images, n=5 independent experiments. GRPR endocytosis is blocked by the clathrin inhibitor PS2 or by the dynamin inhibitor Dy4. **B-F.** GRP activation of GRP recruits βARRs (**B**) and traffics GRPR from the plasma membrane (**C,** KRas-Venus) to Rab5a positive early endosomes (**D, E, F**) in HEK cells. Pretreatment with RC-3095 blocks βARRs recruitment and GRPR endocytosis (**B-F**). Mean±SEM, triplicate technical repeats, n=4-5 independent experiments, **P<0.01, ***P<0.001, ****P<0.0001. One-Way ANOVA, Dunnett’s multiple comparisons.

BRET assays in HEK293 cells quantified the proximity of GRPR-RLuc8 (<10 nm) to expressed protein markers in the plasma membrane (KRas-Venus), early endosomes (Rab5a-Venus) and the recruitment of adapter proteins like the β-arrestins (βAAR1-YFP, βAAR2-YFP). GRP (10 nM) stimulation resulted in the association of GRPR-RLuc8 with βAAR1-YFP and βAAR2-YFP (**Fig. 1 B**) which was blocked by preincubation with the GRPR specific antagonist RC-3095 (10 µM). GRP (10 nM) stimulation resulted in the decreased BRET between GRPR-RLuc8 and the plasma membrane marker KRas-Venus (**Fig. 1 C**) and an increase in the BRET between GRPR-RLuc8 and the early endosomal marker Rab5a-Venus (**Fig. 1 D, E**). Pretreatment with Dy4 or PS2 blocked GRPR removal from the plasma membrane and association with the Rab5a positive early endosomes (**Fig. 1 D, E**). The endocytosis and trafficking to early endosomes was dependent on the activation of GRPR as preincubation with RC-3095 (10 µM) inhibited the increased the GRPR-RLuc8/Rab5a-Venus BRET signal (**Fig. 1 D, E**). Expression of the dominant negative form of dynamin (DynaminK44A) also blocked the trafficking of GRPR from the plasma membrane to early endosomes (**Fig. 1 C, D, E**). Additionally, co-expression of DynK44A significantly inhibited the trafficking of GRPR to Rab7-Venus tagged late endosomes but had no impact on GRPR trafficking to Rab11-Venus tagged recycling endosomes (data not shown).

GRPR persists in an active state and can recruit G proteins in endosomes. The number of GPCRs that can recruit G proteins and signal from endosomes is growing (30). To determine if GRPR could colocalize with G proteins at intracellular compartments, GRPR-mApple was co-expressed with mini (m) Gα proteins in HEK cells. mGα proteins are N-terminally truncated Gα proteins that can diffuse through the cytosol of cells and bind to active conformations of GPCRs (40–42). The mGα_q_ was developed by altering the mGα_s_ residues to mimic the residues and activity of Gα_q_ (41, 42). In HEK293 cells expressing both GRPR-mApple and Venus-mGα_q_, stimulation of GRPR with GRP (10 nM), resulted in the colocalization of Venus-mGα_q_ with GRPR-mApple both at the plasma membrane and at intracellular sites. Confocal micrographs showed that following stimulation with GRP, GRPR-mApple colocalized with Venus-mGα_q_ at intracellular sites (**Fig. 2 B**)

**Figure 2.**
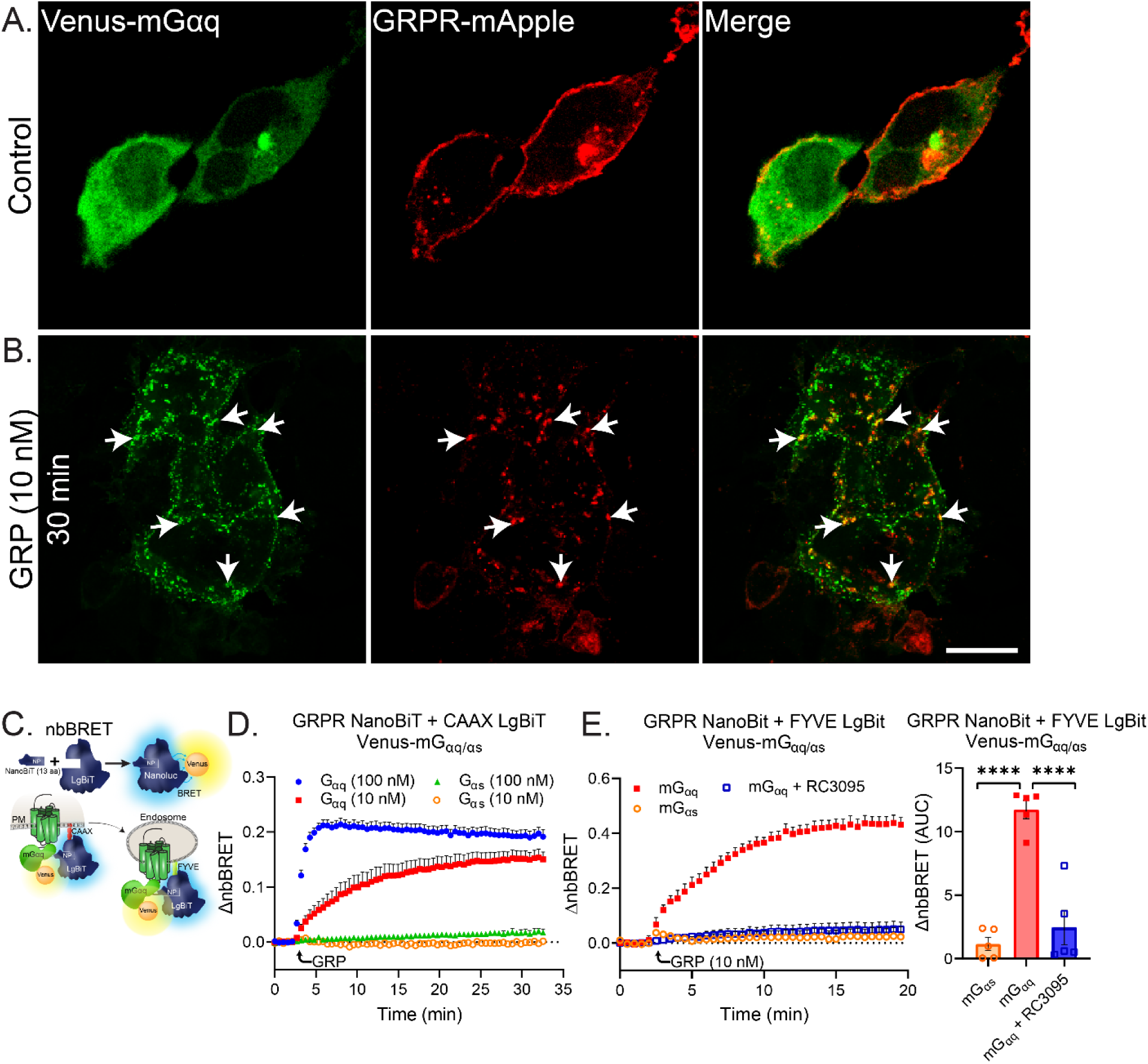
GRPR can recruit miniGα_q_ to early endosomes. A, **B.** HEK293 cells express GRPR-mApple on the plasma membrane while Venus-mGα_q_ is present in the cytosol in quiescent cells. GRP stimulated GRPR internalization and recruitment of Venus-mGα_q_ **(A, B,** Arrows). Scale 25 µm, representative images, n=5 independent experiments. **C**. nbBRET assays used a split nanoluciferase so that when the NP and the LgBiT are in close proximity they combined to form a functional nanoluciferase and when paired with a BRET acceptor, can characterize the colocalization of three distinct proteins. PM = plasma membrane. **D, E**. GRP induced recruitment of Venus-mGα_q/s_ to the plasma membrane (CAAX-LgBiT) in a dose dependent manner (**D**). GRP induced recruitment of Venus-mGα_q/s_ to early endosomes (FYVE-LgBiT) in HEK cells pretreatyed with RC-3095 (**E**). Area under curve (AUC). Mean±SEM, triplicate technical repeats, n=5 independent experiments, ****P<0.0001. One-Way ANOVA, Dunnett’s multiple comparisons.

We used the nanobit-BRET assay (nbBRET) to confirm that GRPR was present in an active state and could associate with G proteins and facilitate intracellular signaling from the endosomal compartments. nbBRET was used to measure the ability of GRPR to recruit and associate with effector proteins, like Gα proteins at different intracellular sites. For the nbBRET assay, NanoLuc luciferase (NLuc) was split into 2 peptides, a small 13 aa NanoBiT fragment (NP) and a larger fragment (LgBiT) (40, 43, 44). When combined, the NP and LgBiT form a functional NanoLuc luciferase that can produce luminescence (∼460 nm wavelength) in the presence of the substrate furimazine (**Fig. 2 C**). We tagged the C-terminal of GRPR with the NP and placed the LgBiT tag on either a plasma membrane marker (CAAX-LgBiT) or an early endosomal marker (FYVE-LgBiT) and expressed them in HEK cells with either Venus-mGα_s_ or Venus-mGα_q_. As GRPR is a Gα_q_ associated GPCR (45), stimulation with GRP drove a dose dependent increase in nbBRET at the plasma membrane between GRPR-NP, CAAX-LgBiT and Venus-mGα_q_ but not with Venus-mGα_s_ (**Fig. 2 D**), showing specificity of the nbBRET assay to detect the recruitment of G proteins to GRPR. To determine if GRPR could recruit Venus-mGα_q_ to endosomes, HEK cells were transfected with the endosomal marker FYVE-LgBiT and challenged with GRP (10 nM) with or without the GRPR antagonist RC-3095 (10 µM). GRP caused a rapid increase in the nbBRET between GRPR-NP, FYVE-LgBiT and Venus-mGα_q_ but not Venus-mGα_s_, showing that GRPR can recruit specific G proteins to the endosomal compartment. Again, mGα_q_ endosomal recruitment was dependent on GRPR activation and was blocked by pretreatment with the antagonist RC-3095 (**Fig. 2 E**).

To investigate how GRPR endocytosis mediates subcellular signaling, we used compartmentalized fluorescence resonance energy transfer (FRET) biosensors. These FRET biosensors can be targeted to subcellular compartments and can provide a kinetic readout to subcellular signaling in HEK cells. We quantified cytosolic (CytoEKAR) and nuclear (NucEKAR) extracellular-regulated kinase activity using FRET biosensors to understand the link between GRPR endocytosis and compartmentalized cellular responses (33, 39). GRP (1 nM) induced a gradual and sustained activation of both cytosolic and nuclear ERK signaling in HEK cells (**Fig. 3**). Pretreatment with the clathrin inhibitor PS2 or the dynamin inhibitor Dyn4 completely blocked both cytosolic (**Fig. 3 A-C**) and nuclear ERK (**Fig. 3 D-F**) signaling, while the negative controls had no impact on signaling. Additionally, the co-expression of DynK44A blocked both cytosolic and nuclear ERK (**Fig. 3 B, C, E, F**) showing the importance of GRPR endocytosis to elicit the complete cellular response to GRP activation.

**Figure 3.**
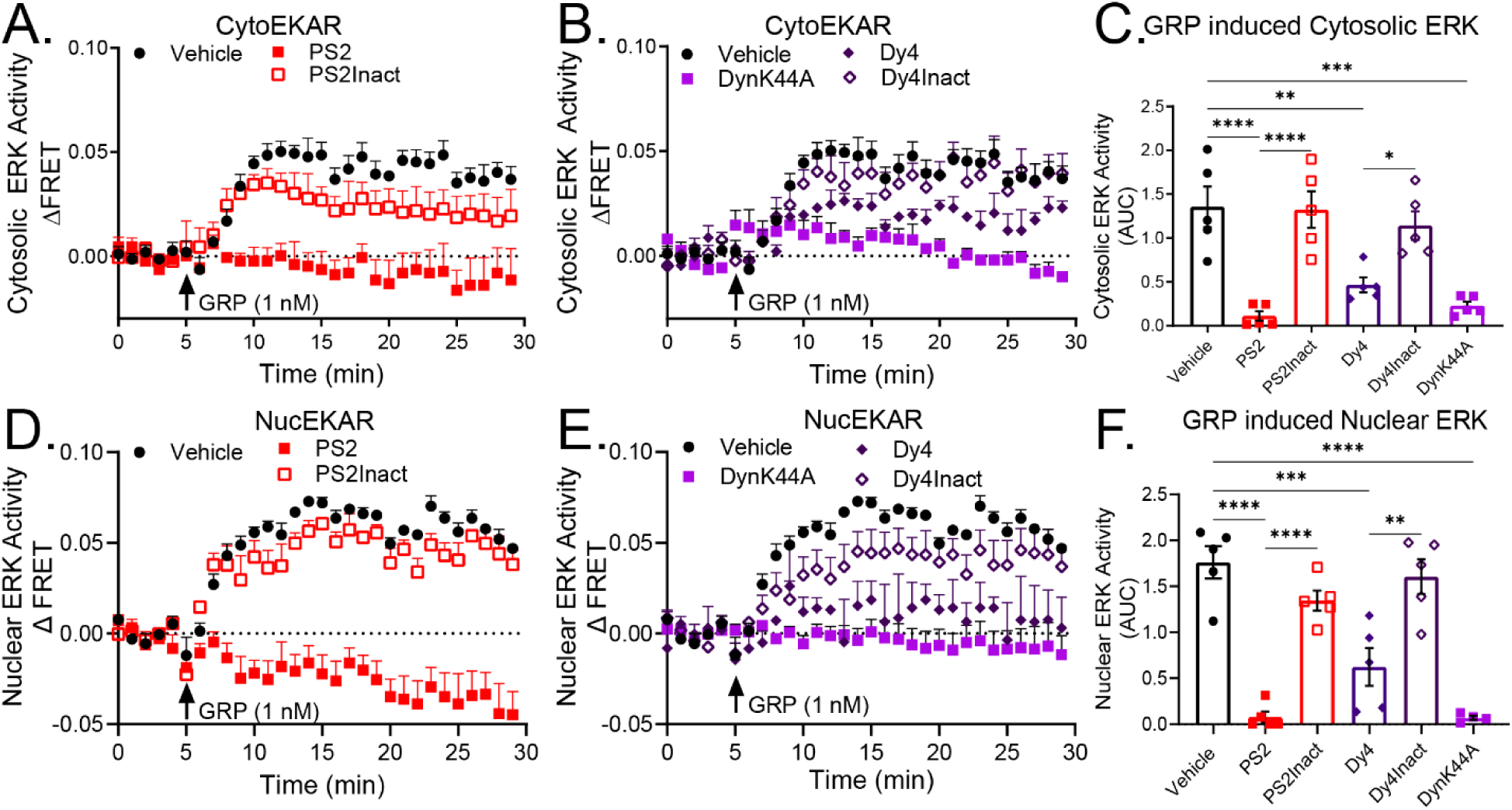
Endosomal signaling of GRPR drives cytosolic and nuclear ERK. GRP induced activation of cytosolic (**A-C**, cytoEKAR) and nuclear (**D-F**, nucEKAR) ERK activation in HEK cells pretreated with PS2 (**A, D**), Dy4 or expressing DynK44A (**B, C**). The negative controls PS2Inact and Dy4Inact had no impact on cytosolic (**C**) or nuclear ERK activation (**F**). Area under curve (AUC). Mean±SEM, triplicate technical repeats, n=5 independent experiments, *P<0.05, **P<0.01, ***P<0.001, ****P<0.0001. One-Way ANOVA, Dunnett’s multiple comparisons.

### Endosomal GRPR signaling mediates itch sensation and scratching

Does the endosomal signaling of GRPR impact itch sensation and scratching behavior in mice? To investigate the role of endosomal GRPR signaling in itch sensation and scratching behavior we administered the endocytic inhibitors PS2 (5 µl/ 30 µM), Dy4 (5 µl/ 30 µM), the inactive controls PS2inact (5 µl/30 µM), Dyn4Inact (5 µl/30µM), or vehicle by intrathecal injection (C4). Mice were allowed to recover for 30 min then mice received an intradermal injection of the pruritogen chloroquine (CQ, 10 µl/ 10 mM) to the nape on the right side. Scratching behavior, grooming, the distance traveled (track length) and inactivity were monitored for 30 min using the behavioral spectrometer (**Fig. 4 A**). Intradermal injection of CQ resulted in an increased number of scratching bouts and total time spent scratching in the vehicle treated mice compared to baseline. Intrathecal injection of PS2 and Dy4 resulted in a significant reduction in both the number of scratching bouts and the time spent scratching during the 30 min observation when both male and female mice were grouped together (**Fig. 4 B**). The inactive compounds PS2Inact and Dy4Inact had no impact on scratching behavior. In male mice, intrathecal injection of PS2 did not significantly inhibit the total time spent scratching (P=0.0668) but trended in an inhibitory manner (**Fig. 4 C**). The number of scratching bouts, however, was significantly inhibited by the intrathecal injection of PS2. Intrathecal Dy4 injections significantly inhibited both the total time spent scratching and the number of scratching bouts in male and female mice when the sexes were analyzed separately (**Fig 4. C, D**). Intrathecal injection of the inactive controls did not impact scratching behaviors in either of the sexes when analyzed separately or in the combined analysis (**Fig. 4 C, D**). The female cohort had a larger behavioral response to CQ, with a greater increase in both scratching bouts and time scratching when compared to the male cohort (**Fig. 4 C, D**). Additionally, pretreatment with PS2 or Dy4 had a more significant reduction in scratching behaviors in the female cohort compared to the male mice (**Fig. 4 C, D**).

**Figure 4.**
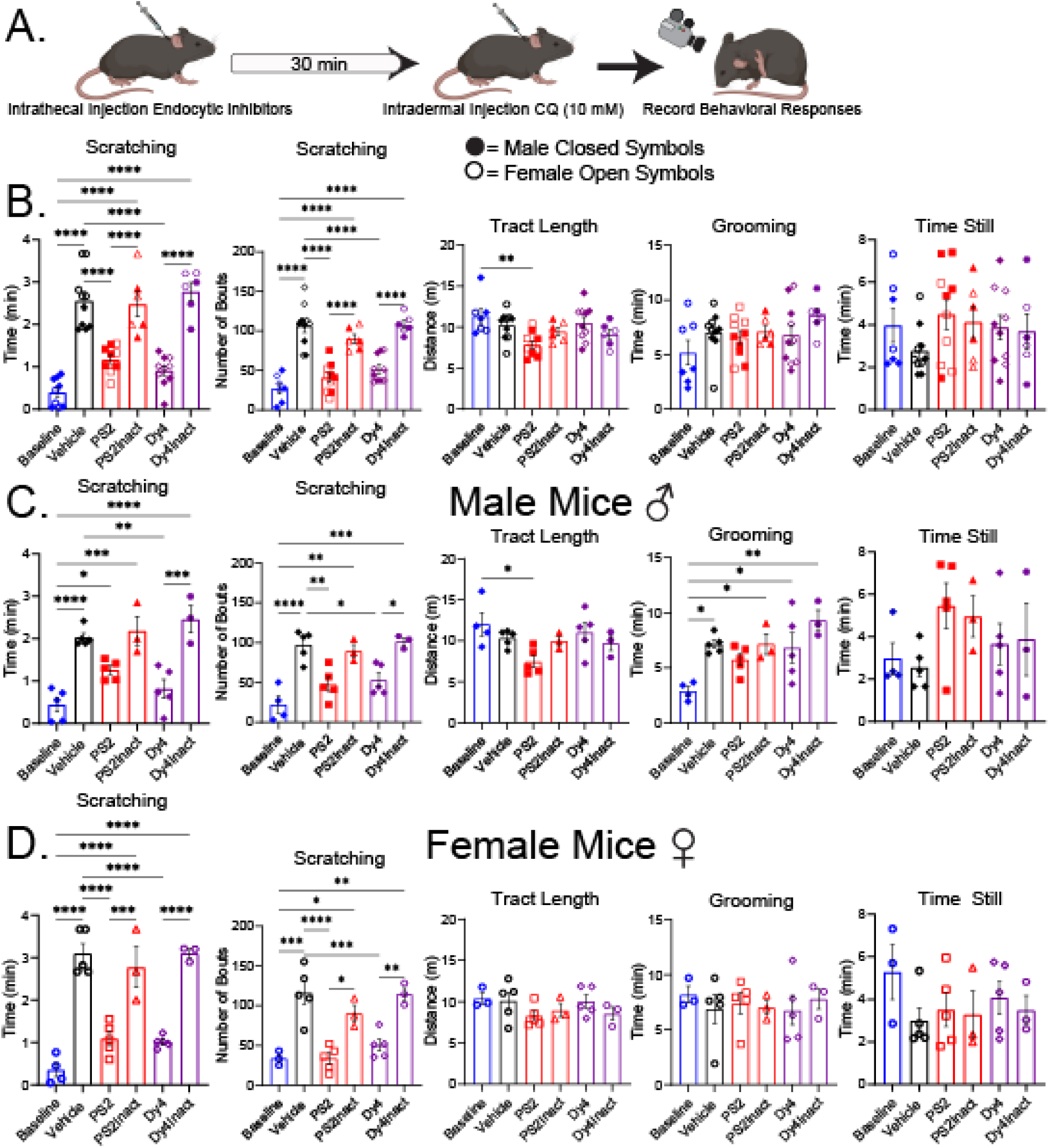
Endocytic inhibitors attenuated CQ induced scratching responses in mice. **A**. Experimental protocol. **B-D**. CQ induced scratching and behavioral patterns in both male and female mice grouped (**B**) and separated to male (**C**) and female (**D**). Endocytic inhibitors block scratching behavior in both sexes without altering normal behavioral patterns. **B** n=6-10 mice, **C**, **D** n=3-5. *P<0.05, **P<0.01, ***P<0.001, ****P<0.0001. Two-way ANOVA, Tukey’s multiple comparison.

Analysis of secondary non-evoked behaviors like tract length, total time spent grooming or total time inactive were not altered by intrathecal injections of either PS2, Dy4, PS2Inact, or Dy4Inact when both sexes were grouped and compared to vehicle treated animals (**Fig. 4 B**). Intrathecal injection of PS2 did result in a small but significant reduction in the tract length during the 30 min observation period compared to the baseline group when male and female mice were analyzed together (**Fig. 4 B**). This reduction in the tract length was only present in the male cohort when compared to the baseline animals (**Fig. 4 C**). Again, there was no significant difference in track length between any of the mice that received the intradermal CQ injection. Intradermal injection of CQ resulted in the increased grooming behavior in male mice compared to baseline and was significant in the vehicle, PS2Inact, Dy4, and Dy4Inact treated groups (**Fig. 4 C**). The increased grooming was only present in the male cohort as grooming behavior in female mice was not altered in any of the groups. Lastly, the time inactive varied greatly between animals and treatment groups and there was no significant difference in the test groups when analyzed together or when mice were separated by sex (**Fig. 4 C, D**).

### siRNA knockdown of dynamin

Pharmacological inhibitors of dynamin are effective at blocking GPCR endocytosis and endosomal signaling (33, 40, 46), but they can have off target effects. Here we used intrathecal injection of siRNA to knockdown dynamin in the spinal cord to confirm the importance of GRPR endocytosis in itch sensation. We have previously used siRNA to target and inhibit dynamin function in the mouse spinal cord in nociceptive assays (33, 34, 47). Using the same technique we used previously (33, 47), we injected a cocktail of siRNA targeting dynamin 1, 2, and 3, or a mismatched control intrathecally (C4) and allowed the mice to recover for 48 h, the point of greatest dynamin knockdown (33, 47). Mice were given intradermal injections of CQ and scratching behaviors were recorded. Similarly to the pharmacologically inhibition of dynamin by Dy4, intrathecal injection of siRNA targeting dynamin significantly reduced scratching behavior in both male and female mice compared to the control mismatched siRNA (**Fig. 5 A**). Dynamin-siRNA did not alter tract length, grooming, or time inactive when both sexes were grouped together (**Fig. 5 A**). However, siRNA targeting dynamin did result in a small but significant reduction in time spent grooming in the female mice but not in the male mice (**Fig. 5 B**). These results, along with the ability of the endocytic inhibitors to block scratching behavior, demonstrate the importance of GRPR internalization and endosomal signaling to activate the GRPR positive interneurons and invoke a scratching response in mice. These studies also highlight the need to target endosomal signaling of GRPR to treat pruritus.

**Figure 5.**
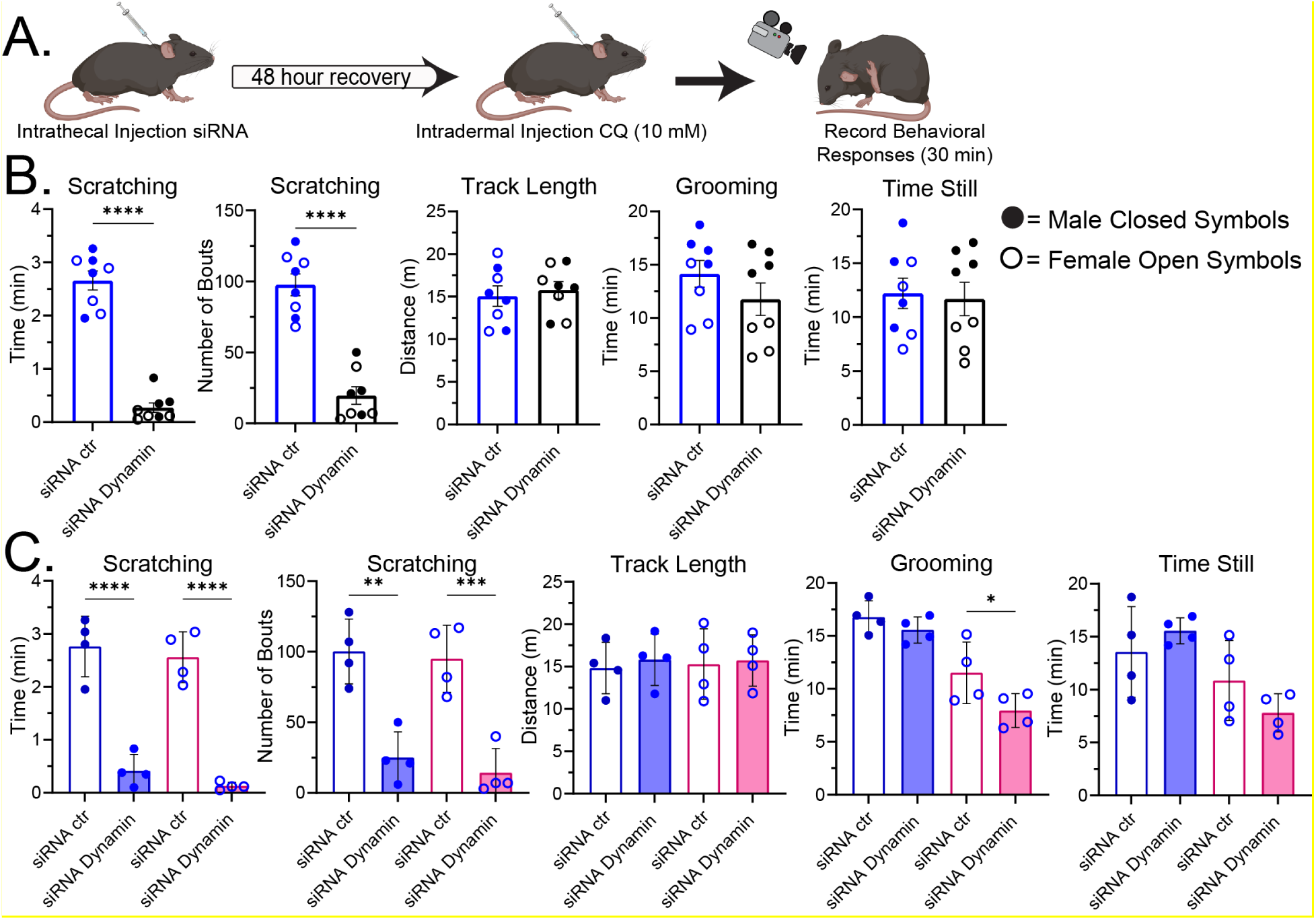
siRNA mediated knockdown of dynamin inhibits CQ induced scratching behavior in mice. **A.** Experimental protocol. **B-C.** Scratching and behavioral responses in mice after intrathecal injection of siRNA targeting dynamin or control mismatched siRNA analyzed as a group (**B**) or separated by sex (**C**). siRNA mediated knockdown of dynamin blocked CQ induced scratching in both male and female mice without altering non-evoked behaviors. **B.** n=8 mice **C.** n=4. *P<0.05, **P<0.01, ***P<0.001, ****P<0.0001. Two-way ANOVA, Tukey’s multiple comparison.

### Targeting endosomal signaling of GRPR

The effectiveness of the endocytic inhibitors and dynamin siRNA to block GRPR trafficking and inhibit signaling pathways in *in vitro* models and itch behavior in mice shows that endosomal signaling of GRPR is essential in activating the itch sensory pathway. We wanted to understand how the delivery of GRPR antagonists to the endosomal compartment would inhibit endosomal GRPR signaling in *in vitro* models and if targeting GRPR antagonists to the intracellular compartment would alter scratching behavior in mice. To deliver the GRPR antagonist to the endosomal compartment of cells we utilized pH-sensitive liposomal coated mesoporous silica nanoparticles.

pH sensitive nanoparticle-encapsulated antagonists inhibit endosomal GRPR signaling. The ability of GRPR to remain in an active state as it traffics and signals from endosomes is a key area of GRPR signaling that has not been explored or targeted. Inhibitors of endocytosis provided a useful tool to understand the role of endocytosis in GRPR signaling but are not a valid therapeutic target as they are not specific to a particular GPCR. Here, we used nanoparticles to deliver GRPR antagonists to specifically target GRPR signaling. We have used pH-responsive nanoparticles to deliver compounds intracellularly and alter endosomal GPCR signaling in previous studies (46, 48). Here, pH-sensitive liposome coated biodegradable mesoporous silica nanoparticles (Lipo-MSNs) were used to deliver the GRPR antagonist RC-3095 intracellularly. Lipo-MSNs were designed to be stable at pH >7.0. However, when the pH is <6.5, as is seen in the endosomes (49), the liposome destabilizes and Lipo-MSN release their cargo (46, 50–52). Lipo-MSNs were spherical nanoparticles with a hydrodynamic diameter ranging between 150-180 nm and had a surface charge of +28 to 36 mv with a polydispersity index of 0.24 to 0.27. Lipo-MSNs had a loading efficiency of RC-3095 of 71 ± 5% and release kinetics were similar to previous studies (46).

To evaluate the ability of Lipo-MSN to target endosomal GRPR signaling, HEK-FLPN-GRPR cells expressing the Rab5a-GFP were incubated with Lipo-MSN-647 for 4 h at 37° C. Cells were washed and challenged with the fluorescently tagged GRP (TAMRA-GRP, 1 µM) for 30 min and imaged using a confocal microscope. Upon addition, Lipo-MSN-647 (40 µg/ml MSN) was bound along the plasma membrane and then internalized and accumulated in the Rab5a-GFP positive endosomes. Following stimulation with TAMRA-GRP, the TAMRA-GRP/GRPR complex internalized and colocalized with Lipo-MSN-647 in the Rab5a-GFP positive endosomes (**Fig. 6 A**).

**Figure 6.**
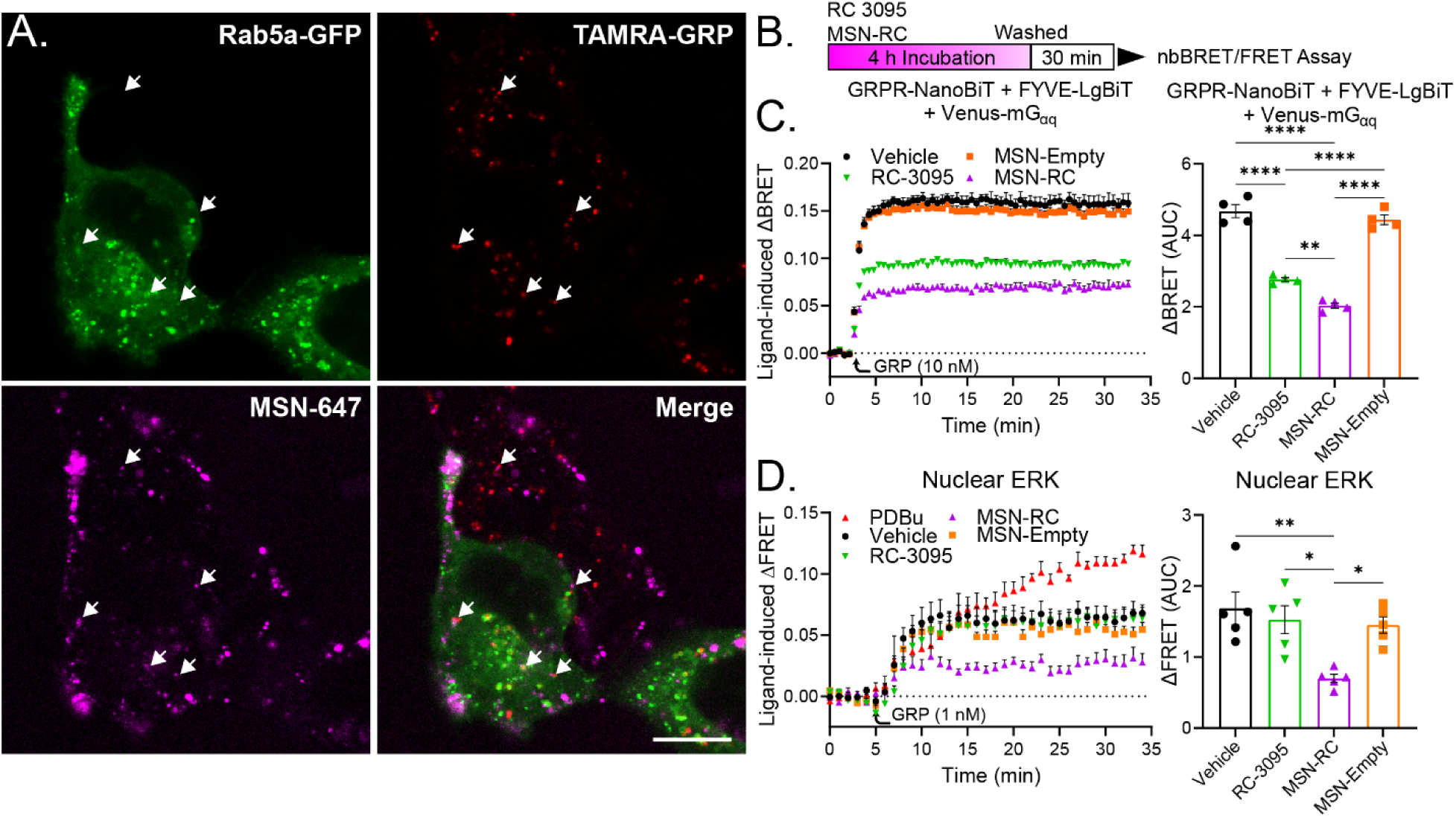
**pH sensitive Lipo-MSN nanoparticles loaded with RC-3095 block Venus-mGα_q_ recruitment and endosomal signaling of GRPR**. **A**. MSN nanoparticles are endocytosed to Rab5a positive early endosomes and colocalized with TAMRA-GRP following a 30 min incubation. Scale 10 µm, representative images, n=4 independent experiments. **B**. Experimental protocol. **C**. nbBRET assay of Venus-mGα_q_ recruitment to GRPR-NP containing early endosomes (FYVE-LgBiT) in HEK293 cells treated with RC-3095 or Lipo-MSNs loaded with RC-3095. MSN-RC showed greater inhibition of miniGα_q_ recruitment (**C**) and nuclear ERK signaling (**D**) compared to free RC. Mean±SEM, triplicate technical repeats, n=4 independent experiments. **D**. GRP stimulated nuclear ERK signaling in HEK-GRPR cells. Mean±SEM, triplicate technical repeats, n=5 independent experiments, *P<0.05, **P<0.01, ****P<0.0001. One-Way ANOVA, Dunnett’s multiple comparisons.

We determined if Lipo-MSN delivery of RC-3095 to the endosomal compartment could inhibit endosomal GRPR signaling. Employing the nbBRET assay used previously, we expressed in the HEK cells GRPR-NP, FYVE-LgBiT, and Venus-mGα_q_ and incubated the cells with Lipo-MSNs loaded with RC-3095 (MSN-RC, 10 µM), empty Lipo-MSNs (MSN-Empty), free RC-3095 (10 µM), or vehicle for 4 h 37 °C. HEK-FLPN-GRPR cells were washed and returned to the incubator for 30 min, then challenged with GRP (10 nM, **Fig. 6 B**). Preincubation with both free RC-3095 and MSN-RC significantly inhibited the endosomal recruitment of Venus-mGα_q._ However, the MSN-RC had a significantly greater inhibition of endosomal recruitment of Venus-mGα_q_ compared to the free RC-3095 (**Fig. 6 C**). MSN-Empty has no impact of Venus-mGα_q_ endosomal recruitment. To further investigate the ability of MSN-RC to inhibit intracellular GRPR signaling, we used the nuclear ERK FRET biosensor assay. HEK-FLPN-GRPR cells expressing the NucEKAR biosensor were incubated with vehicle, RC-3095, MSN-RC or MSN-Empty as in the nbBRET assay (**Fig. 6 B**). HEK-FLPN-GRPR cells were challenged with GRP (1 nM). Only MSN-RC inhibited nuclear ERK signaling (**Fig. 6 D**), while the free RC-3095 and the empty Lipo-MSNs had no effect. These results, paired with the nuclear ERK inhibition of the endocytic inhibitors, demonstrate the importance of intracellular signaling of GRPR to elicit a complete cellular response to GRPR activation and the need to deliver GRPR antagonists intracellularly to block all facets of GRPR signaling.

### Does the intracellular delivery of RC3095 via Lipo-MSN alter scratching behavior in mice?

pH-sensitive Lipo-MSNs are an effective vehicle for the intracellular release of antagonists and compounds (46, 48), but can Lipo-MSNs be used to block endosomal GRPR signaling and inhibit scratching in mice? To test the ability of Lipo-MSNs to be internalized to spinal neurons, mice received intrathecal injection of Lipo-MSNs loaded with the fluorophore Alexa647 (MSN-647). MSN-647 was efficiently endocytosed into the cells in the superficial layers of the dorsal horn of the mouse spinal cord and MSNs loaded with the fluorophore Alexa647 were detected in the dorsal horn at 2 hours following intrathecal injection (**Fig. 7 B**). This is similar to previous studies showing Lipo-MSN uptake into spinal neurons (46, 48). To test the ability of Lipo-MSNs to deliver antagonists intracellular and block endosomal signaling of GRPR, mice received intrathecal injections of either RC-3095 (5 µl/ 10 µM), Lipo-MSN loaded with RC3095 (MSN-RC, 5 µl/ 10 µM, 40 µg/ml), empty Lipo-MSN (MSN-Empty, 5 µl/ 40 µg/ml) or vehicle and allowed to recover for 2 h. Mice then received CQ intradermally (10 µl/ 10 mM) the right nape and behavioral responses were recorded (**Fig. 7 A**). Intrathecal injection of RC-3095 significantly inhibited scratching bouts and time scratching in mice. However, pretreatment with MSN-RC resulted in a more significant inhibition of both time scratching and scratching bouts when compared to the vehicle treated mice and, MSN-RC treatment was more effective at blocking scratching behavior when compared to the free RC-3095 treated mice when analyzing both sexes (**Fig. 7 C**). Empty Lipo-MSN particles had no impact on scratching behavior. MSN-RC nanoparticles had no impact on non-evoked behaviors like tract length, grooming, or time inactive compared to vehicle controls in the 30 min assay when both male and female mice were analyzed together **(Fig. 7 C**). Analysis of the individual sexes showed that both free RC-3095 and MSN-RC significantly inhibited scratching bouts in male and female mice compared to vehicle controls, while only MSN-RC, and not free RC-3095, significantly inhibited scratching time in male mice (**Fig. 7 D**). Together, these data demonstrate that the administration of MSN-RC resulted in a more significant inhibition of scratching time compared to free RC-3095 (**Fig 7. D, E**). While tract length was significantly increased in the free RC-3095 treated mice compared to the MSN-RC treated mice (**Fig. 7 C**), there was no significant change in grooming or time inactive when both sexes were grouped. When analyzed separately, there was no change in tract length or time inactive in either sex (**Fig. 7 D, E**). Female mice showed no change in grooming. However, as observed in earlier experiments, male mice displayed a general increase in grooming behavior following intradermal CQ injections. The increase in grooming was only significant in the RC-3095 and MSN-Empty treatment groups compared to baseline (**Fig. 7 D**).

**Figure 7.**
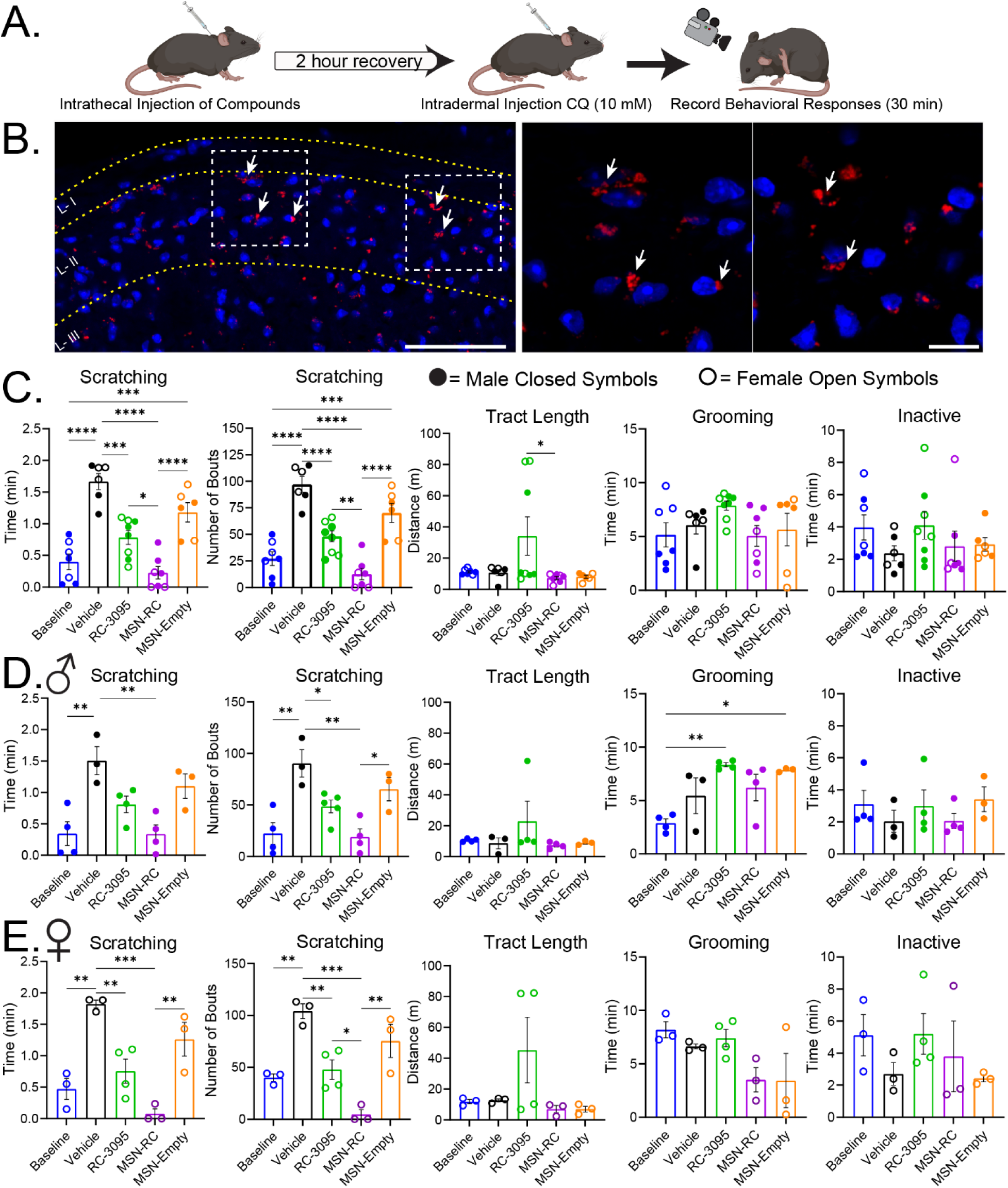
Intrathecal administered MSN-RC inhibits GRPR mediated scratching behaviors in mice. **A**. Experimental protocol. **B.** Lipo-MSNs loaded with Alexa647 are internalized by cells in the dorsal horn of the spinal cord by 2 h post injection. Scale 50 µm and 10 µm for inset. **C-E.** Quantification of CQ induced scratching and secondary behaviors 2 hours after intrathecal injection of vehicle, RC-3095, MSN-RC, or MSN-Empty nanoparticles in mice (**C**). Behavioral responses separated into male (**D**) or female cohorts (**F**). **C.** Mean±SEM n=6-8 mice. **D, E.** n=3-4 mice. *P<0.05, **P<0.01, ***P<0.001, ****P<0.0001. Two-way ANOVA, Tukey’s multiple comparison.

These results demonstrated that pH sensitive Lipo-MSN delivery of a GRPR antagonist can block endosomal signaling of GRPR and more effectively attenuate CQ induced scratching in mice compared to injections of RC-3095 alone. Moreover, these findings support the hypothesis that endosomal signaling of GRPR is a mediator of itch sensation and that the targeting of endosomal GRPR is a viable and more efficacious therapeutic target for treating pruritus.

## Discussion

In this study we have characterized the endosomal trafficking and signaling of the GRPR and that endosomal signaling of GRPR regulates cytosolic and nuclear ERK activation. Moreover, we demonstrate that by inhibiting endocytosis, we can attenuate scratching responses in mice. Our results show that by targeting endosomal GRPR signaling with pH sensitive MSN nanoparticles, we can inhibit endosomal G protein recruitment, block nuclear ERK signaling, and inhibit scratching responses in mouse models for itch. The enhanced efficacy of Lipo-MSN loaded RC-3095 to block scratching responses in mice is likely due to the inhibition of intracellular signaling of GRPR that was evaluated in the *in vitro* assays as free RC-3095 was unable to effectively reach and block GRPR signaling once the receptor was internalized to the endosomal compartment. These studies have characterized a new signaling mechanism for GRPR and have identified a therapeutic approach to deliver antagonists using nanomedicines to block receptor signaling in neurons that regulate itch signaling in the spinal cord.

We used BRET assays to quantify GRPR recruitment of βARRs and G proteins, and to track the movement from plasma membrane to endosomes. FRET based biosensors were used to quantify cytosolic and nuclear ERK activation and to highlight the importance of GRPR trafficking to generate a complete cellular response. These data highlight the ability of endosomal-GRPR to recruit signaling partners like βARRs or miniGα_q_ for prolonged time frames. Indeed, the prolonged endosomal signaling of GRPR would help explain the need for sustained presynaptic release of GRP to stimulate and drive GRPR output in the spinal gate for itch (20). An increasing number of studies have emphasized the importance of prolonged intracellular GPCR signaling events in nociception (33, 40, 46). Those studies together with the data presented here indicate that endosomal trafficking and signaling are necessary for the progression of these pathological states but are less important in driving normal patterns of behavior. A general inhibition of internalization using dynamin or clathrin inhibitors or by dynamin knockdown using siRNA was sufficient to attenuate itch sensation without disrupting general behavioral patterns, recapitulating what has been demonstrated in studies of nociception (39, 44, 48). Only following an elevated level of GPCR activation due to an increase in sensory input, i.e. inflammation or nociception, do we detect an impact of dynamin or clathrin knockdown or inhibition.

Here, we report a similar action in itch sensation at the spinal gate controlled by GRPR (20). As GRP is released in repetitive bursts, more and more GRPR would be internalized and trafficked to endosomes where GRPR would continue to signal and drive the neuronal output necessary for itch sensation. Any extracellular limited antagonist would only target plasma membrane activation of GRPR but would not be able to reach the endosomal network and would miss a large population of endocytosed but active GRPR. Only by targeting the intracellular signaling of GRPR can we fully block the neuronal activation and inhibit itch and scratching in the mouse model. In this study we used an acute model of pruritic where a defined activation of the itch pathway. A study using a chronic model would also be informative as in that model, GRPR would be active for a prolonged time frame over days and weeks and would see an elevated presence in the endosomal network. Hypothetically, MSN encapsulated GRPR antagonists would be even more effective in a chronic model of itch due to the ability of the MSNs to deliver antagonists to the intracellular network where, during chronic activation and prolonged GRP release, more active GRPR would be present and signaling. In this state, an extracellular limited antagonist would not be able to target a large portion of GRPR in these interneurons.

The likelihood of developing chronic itch increases with age and is more prevalent in the female population (53). Here we show an increased scratching response in the female population (**Fig. 4 D, 7 E**). In these cases, MSN-RC was even more effective at blocking scratching responses in the female population compared to the free RC-3095 compound (**Fig. 7 D**). It remains undetermined as to why MSN-RC was more effective at inhibiting scratching in the female population. Is there an increased release of GRP in females or is there a sex dependent difference in the trafficking and regulation of endosomal GRPR signaling? Further studies are needed to better understand the sex-based differences in pruritus and to provide effective therapeutics. This study demonstrates that in order to develop therapeutics that will be effective at treating pruritus, we must consider the ability of GPCRs, like GRPR, to signal from the plasma membrane and from intracellular compartments. Here, pH sensitive nanoparticles have proven to be an effective agent for the intracellular delivery of GRPR antagonists and for the regulation of pruritus.

## Materials and Methods

### Reagents

Human Gastrin releasing peptide (GRP, #24415) and Fura2AM (#14591) were purchased from Cayman Chemicals (Ann Arbor, MI). GRP-TAMRA was produced and purchased from Biomatik (Wilmington, DE). DMEM, Hank’s balanced salt solution (HBSS), and ProLong^TM^ Glass, were purchased from Thermofisher (Waltham, MA). Coelenterazine h was purchased from Nanolight Technology (Pinetop, AZ). PitStop2, PitStop2Inactive, Dyngo4a and Dyngo4aInactive were from Dr. Adam McCluskey (University of Newcastle, AU). All other reagents were from Sigma (St. Louis, MO) unless otherwise specified.

### cDNAs

The pcDNA3.1-hGRPR (GRPR000001) was purchased from cDNA.org. The following plasmids and FRET biosensors were purchased from Addgene: CytoEKAR Cerulean/Venus (Cytosolic ERK sensor, 18679), NucEKAR Cerulean/Venus (Nuclear ERK sensor, 18681), and CytoCKAR CFP/YFP (Cytosolic PKC sensor, 14870), and Dynamin K44A (dominant negative dynamin-1, 34683).

Human Gastrin releasing peptide receptor (GRPR) was cloned into the pcDNA5-FRT vector with or without a C-terminal HA-mApple fluorescent, renilla luciferase 8 (RLuc8) tag, or the NanoBiT (NB) tag using NEBuilder HiFi DNA (New England BioLabs, Ipswich MA). The resulting plasmids were confirmed by sequencing and amplified using chemicompetent DH5α bacteria. KRas-Venus (plasma membrane), Rab5a-Venus (early endosome), Rab7-Venus (late endosomes), Rab11-Venus (recycling endosome), giantin-Venus (cis Golgi body), Venus-miniGα_q_, and Venus-miniGα_s_ were from N. Lambert (Augusta University Medical College of Georgia, GA). βARR2-YFP and βARR1-YFP were from M. Carson, Duke University, Durnham, NC. CAAX-LgBiT (plasma membrane), FYVE-LgBiT (early endosome) were donated by the Alex Thompson lab (New York University).

### Cell Line Characterization and maintenance

HEK-FLPN cells (Thermofisher) were cultured in DMEM media with 10% fetal bovine serum and Zeocin (100µg/ml) and maintained at 37° C and 5% CO_2_. Cells were passaged at 90% confluency.

### Stable HEK-GRPR and HEK-GRPR-mApple cell line

To make stable GRPR and GRPR-mApple expressing cells, HEK-FLPN cells were plated in a 10 cm dish and transfected with pcDNA5FRT_GRPR or pcDNA5-FRT_GRPR-mApple (1 µg) with the recombinase pOG44 (5 µg) using the PEI transfection method with a ratio of DNA:PEI of 1:6. GRPR or GRPR-mApple stable expressing cells were selected using hygromycin (100 µg/ml) resistance for 5 days. Single colonies surviving selection were collected and maintained DMEM with 10% FBS and hygromycin (100 µg/ml). Functional activity of GRPR and GRPR-mApple in HEK cells were confirmed via Ca^2+^ signaling in response to GRP. A single colony of each cell line was then amplified and used for the following experiments.

### In Vitro Assays

#### GRPR Internalization

HEK-GRPR-mApple or HEK-FLPN-GRPR cells were plated on PDL coated 12 mm cover glass in a 24 well plate or on 35 mm glass bottom dishes (MatTek, Ashland MA). Cells were transduced with the CellLight Early Endosome-GFP (Thermofisher) to express Rab5a-GFP, an early endosomal marker. Cells were incubated overnight. Cells were washed and placed in Hanks balanced salt solution (HBSS, +10 mM HEPES pH 7.4) and incubated with the endocytic inhibitors PitStop (PS, 30 µM), Dyngo4a (Dy4, 30 µM), vehicle or the inactive controls PitNot (PSInact, 30 µM) or Dyngo4a Inactive (Dy4Inact, 30 µM) for 1 h at 37° C. Glass bottom dishes were moved to the microscope for live cell imaging. Cells were challenged with GRP or TAMRA-GRP (10 nM) and recorded for 30 min with the Leica SP8 confocal microscope. HEK-GRPR-mApple cells cultured on 12 mm coverslips were challenged with GRP (10 nM) for 0, 5, 20, or 30 minutes at 37° C. The 12 mm coverslips were fixed in PBS + 4% paraformaldehyde for 20 min on ice and washed with 3x in PBS. To visualize Venus-miniGα_q_ recruitment to GRPR, GRPR-mApple cells were transfected with Venus-miniGα_q_ (3 µg) using the PEI method. Cells were challenged with vehicle or GRP (10 nM) for 30 min and fixed with 4% paraformaldehyde. Cover slips were mounted on glass slides with ProLong Glass mounting medium (Thermofisher). Cells were imaged with a Leica SP8 confocal with a 40x or 63x (NA=1.42) objective. Micrographs were analyzed in FIJI (ImageJ, NIH) and figures were made in Adobe Illustrator.

#### GRPR BRET and nbBRET assays

To characterize the translocation of GRPR from the plasma membrane to the early and late endosomal compartments, HEK293 were transfected in 10 cm dishes with the following plasmids: GRPR-RLuc8 (1 µg) and KRas-Venus (3 µg), Rab5a-Venus (3 µg), Rab7-Venus (3 µg), Rab11-Venus (3 µg), giantin-Venus (3 µg) βARR1-YFP or βARR2-YFP (3 µg) with or without a co-transfection of DynK44A (1 µg). To characterize the recruitment of G proteins to the plasma membrane or to endosomes, HEK293 cells were transfected with GRPR-NanoBiT (1 µg) and CAAX-LgBiT (2 µg) + Venus-miniGα_q/s_ (2 µg) or FYVE-LgBiT (2 µg) + Venus-miniGα_q/s_ (2 µg) using the PEI method with a DNA:PEI ratio of 6:1. Cells were plated in PDL coated 96 well white walled plates and incubated overnight in normal growth media. Cells were washed 2x in HBSS and incubated with the endocytic inhibitors PS, Dy4 (30 µm), inactive controls PSInact or Dy4Inact (30 µM), the GRPR antagonist RC-3095 (10 µM), or vehicle for 1 h. The cells were incubated in the substrate for RLuc8 (coelenterazine H, 2.5 µM) or Nano luciferase (furimazine, 10 µM) for 15 min. BRET and nbBRET was quantified for 30 min using the CLARIOStar plate reader (BMG Labtech, Cary NC) using BRET and nbBRET filters (donor filter: 460 ± 40 nm, acceptor filter: 540 ± 25 nm). Baseline was recorded for 2.5-5 min and GRP (10 nM to 1 µM) was added to the cells. The BRET signal was calculated as the ratio of the acceptor (YFP or Venus) emission over the donor (RLuc8, Nanoluc) emission. Data is normalized to the baseline BRET or nbBRET values and to vehicle treated cells.

#### GRPR FRET assays

FRET biosensors were used to assess subcellular compartmentalized signaling of GRPR. HEK-FLPN-GRPR cells were transfected with CytoCKAR (4 µg), CytoEKAR (4 µg), or NucEKAR (4 µg) FRET biosensors with or without DynaminK44A (1 µg) using the PEI method. Cells were plated in PDL coated black clear bottom plates in serum free media and incubated overnight. Cells were washed 2x in HBSS and incubated with the endocytic inhibitors PS, Dy4, inactive controls PSInact or Dy4Inact (30 µM), RC-3095 (10 µM), or vehicle for 1 h. Cells were placed in the CLARIOstar plate reader. Baseline FRET was recorded for 5 min and the cells were challenged with GRP (1-10 nM). Changes in the FRET ratio were recorded and normalized to the baseline and to vehicle treated cells.

### Endosomally sensitive Mesoporous Nanoparticle (MSNs) Synthesis

#### Liposome and MSN Generation

The biodegradable MSNs were synthesized by the modified sol-gel method with cetyltrimethylammonium tosylate (CTAT) as a template and tetraethyl orthosilicate (TEOS) and bis-silylated precursors with diselenide-bridged groups as co-silica sources (51). Self-assembling liposomes were made utilizing the thin-film hydration method with 1,2-dioleoyl-3-trimethylammonium propane (DOTAP), cholesterol, dioleoyl-phosphatidyl-ethanolamine (DOPE), and DSPE-PEG2000 in a 65:30:3.75:12.5 molar mass ratio in chloroform. Briefly, DOTAP, cholesterol, DOPE, and half of DSPE-PEG2000 were combined in a chloroform solution. The chloroform was evaporated under vacuum and dried overnight. The film was hydrated using a solution of ethanol/sodium acetate buffer (200mM, pH 5.2, vol:vol, 9:1). The film solution was added dropwise to an aqueous solution containing the other half of DSPE-PEG and mixed for 15 minutes. The solution was then incubated at 37° C for 30 minutes and then purified by dialysis against water in 3500 molecular weight cut-off cassettes (54).

#### MSN core-loading and liposome coating

The GRPR antagonist RC-3095 and/or Alexa647 was encapsulated inside the MSN to examine GRPR endosomal signaling. RC-3095 and/or Alexa647 was added to aqueous MSN in 1:10 ratio (wt:wt) and was encapsulated in the MSN through shaking at 750 rpm at 4° C for 30 minutes. Excess cargo was removed by centrifugation at 20,000g, 4° C for 15 minutes. The supernatant was removed, and pellets were resuspended by pipetting in water. The amount of RC-3095 or Alexa647 loaded was quantified by calculating the drug concentrations before and after encapsulation by MSNs. For liposome coating of drug loaded MSNs, the MSN solution was mixed with liposomes in a 1:1 ratio (wt:wt) and shaken at 750 rpm at 4° C for 30 minutes. Excess cargo and liposomes were removed by centrifugation at 20,000g, 4° C for 15 minutes and the LipoMSN-RC pellet was resuspended in water and sonicated for 2 minutes.

#### Characterization of MSNs

The hydrodynamic diameter and zeta potential of the NPs in water or PBS were characterized using a Nano-ZS 90 Nanosizer (Malvern Instruments Ltd., Worcestershire, UK).

#### Endosomal trafficking of MSNs

HEK-FLPN-GRPR cells transduced with the CellLight Early Endosome marker were incubated with MSN-647 (40 µg/ml) for 4 hours to characterize cellular uptake as done in previous studies with HEK cells and MSN nanoparticles (46). At the 4 h time point, cells were washed with HBSS and allowed to recover for 30 min. HEK-FLPN-GRPR cells were challenged with TAMRA-GRP (1 µM, 30 min). Live cell imaging followed TAMRA-GRP/GRPR activation and endocytosis to early endosomes. Colocalization of MSN-647 in early endosomes (Rab5a-GFP) with TAMRA-GRP was assessed HEK-FLPN-GRPR cells, 30 min after TAMRA-GRP addition.

### MSN-RC3095 and GRPR signaling

<u>nbBRET assays</u>. HEK293 cells were transfected with GRPR-NanoBiT (1 µg), the plasma membrane marker CAAX-LgBiT (1 µg), and Venus-miniGα_q_ (3 µg) or Venus-miniGα_s_ (3 µg) or GRPR-SmBiT (1 µg), the endosomal marker FYVE-LgBiT (1 µg), and Venus-miniGα_q_ (3 µg) or Venus-miniGα_s_ (3 µg) as described above. Cells were plated in white walled 96 well plates as described. Cells were incubated with vehicle, RC-3095 (10 µM), MSN particles loaded with RC-3095 (MSN-RC, 40 µg/ml of Lipo-MSN, 10 µM of RC-3095), or empty MSN nanoparticles (MSN-Empty, 40 µg/ml) for 4 h and washed in HBSS 2x. Cells were incubated for 30 min at 37° C, then challenged with GRP (10 nM) and the BRET responses were recorded with the CLARIOstar plate reader.

#### Nuclear ERK assays

HEK-GRPR cells were transfected with the NucEKAR (4 µg) FRET biosensor and incubated for 24 h. Cells were plated in a black walled 96 well plate (30,000 cells/well) and serum starved overnight. Cells were incubated for 4 h with vehicle, RC-3095 (10 µM), MSN-RC3095 (40 µg/ml, 10 µM) or empty MSN nanoparticles. Cells were washed 2x in HBSS and allowed to recover for 30 min. Cells were then challenged with GRP (1 nM) and FRET responses recorded.

### Animal Studies

The institutional animal care and use committee of New York University approved all protocols used in this study.

### Animals

Male and female C57BL/6J mice (8 weeks old) were purchased from Jackson research laboratories (#00064 Jax®, wild-type) and were maintained in a temperature-controlled environment (22° C), on a 12-h light/dark cycle and with free access to food and water. Mice were killed by anesthetic (isoflurane) overdose and by bilateral thoracotomy.

### MSN-Alexa647 uptake in the spinal cord

Mice were lightly anesthetized (isoflurane) and shaved along the neck. MSN loaded with Alexa647 fluorophore were injected intrathecally (5 µl, 40 µg/ml) into the cervical region of each mouse. Mice were returned to their home cage and allowed to recover for 2 h. Mice were then anesthetized and sacrificed (Isoflurane), perfused with ice cold PBS, and 4% paraformaldehyde and the spinal cord was processed as stated earlier. The spinal cord was sectioned (30 µm) and placed on Superfrost slides, stained with DAPI (500 nM) and mounted with ProLong Glass mountant. Sections were imaged on the Leica SP8 confocal to determine the uptake and retention of MSN-Alexa647 in the spinal sections.

### siRNA mediated knockdown of dynamin

siRNA mediated knockdown of dynamin was performed as previously described (33, 40, 55). Briefly, a cationic liposome and adjuvant ionic polymer (polyglutamate) were used to intrathecally deliver ON-TARGETplus siRNA (Dharmacon) targeting mouse dynamin-1, 3 or control siRNA. siRNA (50 ng, 0.5 µl of 100 ng/µl stock) was mixed with 0.5 µl of the adjuvant polyglutamate solution (0.1 µg/µl stock) and 1.5 µl of a sterile 0.15 M NaCl solution. Liposome solution, cationic lipid 2-{3-[bis-(3-amino-propyl)-amino]-propylamino}-N-ditetradecylcarbamoylmethyl-acetamide (DMAPAP) and L-α-dioleoyl phosphatidylethanolamine (DOPE) (DMAPAP/DOPE, 1/1 M:M) (2.5 μl of 200 μM) was added to the siRNA/adjuvant mixture, vortexed for 1 min, and incubated for 30 min at room temperature (33). The siRNA lipoplexes were administered to mice by intrathecal injection (C4, 5 μl). Behavioral tests were performed 48 h after siRNA injection.

### Itch Behavioral studies

To assess endosomal inhibitors on pruriception in the skin, mice from both sexes were lightly anesthetized with isoflurane and shaved along the nape of the neck on the right side. Mice received an intrathecal injection (C4, 5 µl) of PS2 (30 µM), Dy4 (30 µM), PSInact (30 µM), Dy4Inact (30 µM), RC-3095 (10 µM) or vehicle in saline. Mice were allowed to recover for 30 min then chloroquine (10 µl of 10 mM in saline) was injected intradermally into the nape of the neck on the right side. Mice were placed in the Behavioral Spectrometer (Biobserve, Wilhelmstr Germany) and recorded for 30 min. Scratching bouts, time scratching, time grooming, time motionless, and distance traveled was recorded for each mouse.

To assess the effectiveness of MSN nanoparticle mediated delivery of GRPR antagonists to block scratching in mice, mice were prepared as stated but received a 5 µl intrathecal injection of RC-3095 (10 µM), vehicle, MSN-RC (40 µg/ml, 10 µM) or MSN-Empty (40 µg/ml) in sterile saline. Mice were allowed to recover for 2 h in their home cage. Mice were given an intradermal injection to the nape of the neck of chloroquine (10 µl, 10 mM in saline) and recorded for 30 min in the behavioral spectrometer.

### Data analysis and statistics

Sample size for the experiments was determined by a power analysis of our previous studies and was not altered during the investigation. *In vitro* experiments using HEK293 cells were replicated in ≥5 separate experiments, each with 3 technical replicates for each group. The duration of the experiments was determined prior to the beginning of the study. Cells were excluded from analysis if they were contaminated. Experiments using mice were replicated in >8 animals per group with >4 animals of the female or male sex. All mice were of the same age, body weight, and in good health. The protocol for stopping any *in vivo* experiment was defined prior to the beginning of the study. All collected data during this study were included in the manuscript. Outliers were not removed from the experiments. Results were analyzed using Microsoft Excel and Prism 10 and graphs were made using Prism 10. Data sets were analyzed for statistical significance using one-way ANOVA with a Dunnett’s *post hoc* test or a two-way ANOVA with a Tukey’s *post hoc* test. Results are expressed as mean±SEM. The level of significance was set at P ≤ 0.05.

## Acknowledgements

N.A. Lambert (Augusta University) provided cDNA encoding mGα isoforms coupled to Venus. A. Thomsen (New York University) provided cDNA encoding CAAX and FYVE. We thank Tracy Chiu and Evan Chen for technical assistance. This research was funded by NIH R01NS125413 (DDJ).

## References

1. G. Yosipovitch, J. D. Rosen, T. Hashimoto, Itch: From mechanism to (novel) therapeutic approaches. J Allergy Clin Immunol 142, 1375–1390 (2018).

2. X. J. Chen, Y. G. Sun, Central circuit mechanisms of itch. Nat Commun 11, 3052 (2020).

3. Z. F. Chen, A neuropeptide code for itch. Nat Rev Neurosci 22, 758–776 (2021).

4. T. Akiyama et al., Roles for substance P and gastrin-releasing peptide as neurotransmitters released by primary afferent pruriceptors. J Neurophysiol 109, 742–748 (2013).

5. T. Akiyama, M. Tominaga, K. Takamori, M. I. Carstens, E. Carstens, Roles of glutamate, substance P, and gastrin-releasing peptide as spinal neurotransmitters of histaminergic and nonhistaminergic itch. Pain 155, 80–92 (2014).

6. T. Buhl et al., Protease-Activated Receptor-2 Regulates Neuro-Epidermal Communication in Atopic Dermatitis. Front Immunol 11, 1740 (2020).

7. F. Cevikbas et al., A sensory neuron-expressed IL-31 receptor mediates T helper cell-dependent itch: Involvement of TRPV1 and TRPA1. J Allergy Clin Immunol 133, 448–460 (2014).

8. K. Sakai et al., Mouse model of imiquimod-induced psoriatic itch. Pain 157, 2536–2543 (2016).

9. T. Akiyama, M. I. Carstens, E. Carstens, Facial injections of pruritogens and algogens excite partly overlapping populations of primary and second-order trigeminal neurons in mice. J Neurophysiol 104, 2442–2450 (2010).

10. T. Lieu et al., The bile acid receptor TGR5 activates the TRPA1 channel to induce itch in mice. Gastroenterology 147, 1417–1428 (2014).

11. K. Tsujii, T. Andoh, J. B. Lee, Y. Kuraishi, Activation of proteinase-activated receptors induces itch-associated response through histamine-dependent and -independent pathways in mice. J Pharmacol Sci 108, 385–388 (2008).

12. Q. Liu et al., Sensory neuron-specific GPCR Mrgprs are itch receptors mediating chloroquine-induced pruritus. Cell 139, 1353–1365 (2009).

13. D. Pavlenko et al., IL-23 modulates histamine-evoked itch and responses of pruriceptors in mice. Exp Dermatol 29, 1209–1215 (2020).

14. J. Meixiong, C. Vasavda, S. H. Snyder, X. Dong, MRGPRX4 is a G protein-coupled receptor activated by bile acids that may contribute to cholestatic pruritus. Proc Natl Acad Sci U S A 116, 10525–10530 (2019).

15. T. Akiyama, E. Carstens, Neural processing of itch. Neuroscience 250, 697–714 (2013).

16. X. Dong, X. Dong, Peripheral and Central Mechanisms of Itch. Neuron 98, 482–494 (2018).

17. S. Sun, X. Dong, Trp channels and itch. Semin Immunopathol 38, 293–307 (2016).

18. C. Moore, R. Gupta, S. E. Jordt, Y. Chen, W. B. Liedtke, Regulation of Pain and Itch by TRP Channels. Neurosci Bull 34, 120–142 (2018).

19. S. R. Wilson et al., The ion channel TRPA1 is required for chronic itch. J Neurosci 33, 9283–9294 (2013).

20. M. Pagani et al., How Gastrin-Releasing Peptide Opens the Spinal Gate for Itch. Neuron 103, 102–117 (2019).

21. B. Aresh et al., Spinal cord interneurons expressing the gastrin-releasing peptide receptor convey itch through VGLUT2-mediated signaling. Pain 158, 945–961 (2017).

22. T. D. Sheahan, C. A. Warwick, L. G. Fanien, S. E. Ross, The Neurokinin-1 Receptor is Expressed with Gastrin-Releasing Peptide Receptor in Spinal Interneurons and Modulates Itch. J Neurosci 40, 8816–8830 (2020).

23. E. Polgar et al., Grpr expression defines a population of superficial dorsal horn vertical cells that have a role in both itch and pain. Pain 164, 149–170 (2023).

24. X. Liu et al., GRPR/Extracellular Signal-Regulated Kinase and NPRA/Extracellular Signal-Regulated Kinase Signaling Pathways Play a Critical Role in Spinal Transmission of Chronic Itch. J Invest Dermatol 10.1016/j.jid.2020.09.008 (2020).

25. Y.-G. Sun, Z.-F. Chen, A gastrin-releasing peptide receptor mediates the itch sensation in the spinal cord. Nature 448, 700 (2007).

26. Y. G. Sun et al., Cellular basis of itch sensation. Science 325, 1531–1534 (2009).

27. C. Forster, H. O. Handwerker, “Central Nervous Processing of Itch and Pain” in Itch: Mechanisms and Treatment, E. Carstens, T. Akiyama, Eds. (Boca Raton (FL), 2014).

28. G. S. Kroog, X. Jian, L. Chen, J. K. Northup, J. F. Battey, Phosphorylation uncouples the gastrin-releasing peptide receptor from G(q). J Biol Chem 274, 36700–36706 (1999).

29. P. J. Donohue et al., An aspartate residue at the extracellular boundary of TMII and an arginine residue in TMVII of the gastrin-releasing peptide receptor interact to facilitate heterotrimeric G protein coupling. Biochemistry 38, 9366–9372 (1999).

30. N. G. Tsvetanova, R. Irannejad, M. von Zastrow, G protein-coupled receptor (GPCR) signaling via heterotrimeric G proteins from endosomes. J Biol Chem 290, 6689–6696 (2015).

31. E. F. Grady et al., Direct observation of endocytosis of gastrin releasing peptide and its receptor. J Biol Chem 270, 4603–4611 (1995).

32. S. Rajagopal, S. K. Shenoy, GPCR desensitization: Acute and prolonged phases. Cell Signal 41, 9–16 (2018).

33. D. D. Jensen et al., Neurokinin 1 receptor signaling in endosomes mediates sustained nociception and is a viable therapeutic target for prolonged pain relief. Sci Transl Med 9 (2017).

34. R. E. Yarwood et al., Endosomal signaling of the receptor for calcitonin gene-related peptide mediates pain transmission. Proc National Acad Sci 114, 12309–12314 (2017).

35. N. G. Tsvetanova, M. von Zastrow, Spatial encoding of cyclic AMP signaling specificity by GPCR endocytosis. Nature Chemical Biology 10.1038/nchembio.1665, 1–6 (2014).

36. J. E. Murphy, B. E. Padilla, B. Hasdemir, G. S. Cottrell, N. W. Bunnett, Endosomes: a legitimate platform for the signaling train. Proc Natl Acad Sci U S A 106, 17615–17622 (2009).

37. Alex R. B. Thomsen et al., GPCR-G Protein-β-Arrestin Super-Complex Mediates Sustained G Protein Signaling. Cell 166, 907–919 (2016).

38. Y. Kwon et al., Non-canonical beta-adrenergic activation of ERK at endosomes. Nature 611, 173–179 (2022).

39. N. N. Jimenez-Vargas et al., Protease-activated receptor-2 in endosomes signals persistent pain of irritable bowel syndrome. Proc Natl Acad Sci U S A 115, E7438–E7447 (2018).

40. R. Latorre et al., Mice expressing fluorescent PAR(2) reveal that endocytosis mediates colonic inflammation and pain. Proc Natl Acad Sci U S A 119 (2022).

41. Q. Wan et al., Mini G protein probes for active G protein-coupled receptors (GPCRs) in live cells. J Biol Chem 293, 7466–7473 (2018).

42. R. Nehme et al., Mini-G proteins: Novel tools for studying GPCRs in their active conformation. PLoS One 12, e0175642 (2017).

43. A. S. Dixon et al., NanoLuc Complementation Reporter Optimized for Accurate Measurement of Protein Interactions in Cells. ACS Chem Biol 11, 400–408 (2016).

44. A. Hegron et al., Therapeutic antagonism of the neurokinin 1 receptor in endosomes provides sustained pain relief. Proc Natl Acad Sci U S A 120, e2220979120 (2023).

45. M. R. Hellmich, J. F. Battey, J. K. Northup, Selective reconstitution of gastrin-releasing peptide receptor with G alpha q. Proc Natl Acad Sci U S A 94, 751–756 (1997).

46. N. N. Jimenez-Vargas et al., Endosomal signaling of delta opioid receptors is an endogenous mechanism and therapeutic target for relief from inflammatory pain. Proc Natl Acad Sci U S A 117, 15281–15292 (2020).

47. R. Tonello et al., The contribution of endocytosis to sensitization of nociceptors and synaptic transmission in nociceptive circuits. Pain 164, 1355–1374 (2023).

48. P. D. Ramirez-Garcia et al., A pH-responsive nanoparticle targets the neurokinin 1 receptor in endosomes to prevent chronic pain. Nat Nanotechnol 14, 1150–1159 (2019).

49. C. Wang et al., Investigation of endosome and lysosome biology by ultra pH-sensitive nanoprobes. Adv Drug Deliv Rev 113, 87–96 (2017).

50. L. Xiong et al., Tunable stellate mesoporous silica nanoparticles for intracellular drug delivery. J Mater Chem B 3, 1712–1721 (2015).

51. D. Shao et al., Bioinspired Diselenide-Bridged Mesoporous Silica Nanoparticles for Dual-Responsive Protein Delivery. Adv Mater 10.1002/adma.201801198, e1801198 (2018).

52. C. Pinese et al., Sustained delivery of siRNA/mesoporous silica nanoparticle complexes from nanofiber scaffolds for long-term gene silencing. Acta Biomater 76, 164–177 (2018).

53. U. Matterne, C. J. Apfelbacher, L. Vogelgsang, A. Loerbroks, E. Weisshaar, Incidence and determinants of chronic pruritus: a population-based cohort study. Acta Derm Venereol 93, 532–537 (2013).

54. M. Wang et al., Efficient delivery of genome-editing proteins using bioreducible lipid nanoparticles. Proc Natl Acad Sci U S A 113, 2868–2873 (2016).

55. A. Schlegel et al., Anionic polymers for decreased toxicity and enhanced in vivo delivery of siRNA complexed with cationic liposomes. J Control Release 152, 393–401 (2011).

